# Characterization of red fluorescent reporters for dual-color *in vivo* three-photon microscopy

**DOI:** 10.1101/2021.10.18.464840

**Authors:** Michael A. Thornton, Gregory L. Futia, Michael E. Stockton, Baris N. Ozbay, Karl Kilborn, Diego Restrepo, Emily A. Gibson, Ethan G. Hughes

## Abstract

**Significance:** Three-photon (3P) microscopy significantly increases the depth and resolution of *in vivo* imaging due to decreased scattering and nonlinear optical sectioning. Simultaneous excitation of multiple fluorescent proteins is essential to study multicellular interactions and dynamics in the intact brain.

**Aim:** We characterized the excitation laser pulses at a range of wavelengths for 3P microscopy, and then explored the application of tdTomato or mScarlet and EGFP for dual-color single-excitation structural 3P imaging deep in the living mouse brain.

**Approach:** We used frequency-resolved optical gating to measure the spectral intensity, phase, and retrieved pulse widths at a range of wavelengths. Then, we performed *in vivo* single wavelength-excitation 3P imaging in the 1225 - 1360 nm range deep in the mouse cerebral cortex to evaluate the performance of tdTomato or mScarlet in combination with EGFP.

**Results:** We find that tdTomato and mScarlet, expressed in oligodendrocytes and neurons respectively, have high signal-to-background in the 1300–1360 nm range, consistent with enhanced 3P cross sections.

**Conclusions:** These results suggest that a single excitation wavelength source is advantageous for multiple applications of dual-color brain imaging and highlight the importance of empirical characterization of individual fluorophores for 3P microscopy.

## 1. Introduction

Multiphoton microscopy methods allow for high-resolution structural and functional imaging deep into biological tissues^1–3^. Two-photon (2P) microscopy is a widely used technique to assess cellular and subcellular dynamics in the intact mouse brain. Although multiple excitation source 2P microscopy setups are increasingly common^4,5^, broad excitation profiles of specific green and red fluorescent proteins allow simultaneous dual-color 2P imaging using a single tunable excitation source^6^. This single-wavelength excitation dual-color 2P microscopy approach has benefited from careful characterizations of fluorescent protein excitation cross sections at relevant wavelengths^7–9^ and enabled both functional and structural interrogation of neural dynamics *in vivo*. For example, simultaneous 1000 nm excitation of GCaMP6s / jRGECO1a permitted the detection of correlated axonal and dendritic calcium events in the visual cortex^10^, while 920 nm excitation of EGFP / tdTomato was used to track oligodendroglial differentiation and cell fate over time^11^. However, *in vivo* dual color 2P microscopy is practically limited in imaging depth to the superficial mouse cortex by light scattering and the generation of out-of-focus background.

The development of three-photon (3P) microscopy, which utilizes low rep-rate, high pulse energy excitation sources at 1300 - 1700 nm, has significantly improved imaging depth and resolution due to decreased scattering and improved signal to background ratio^3,12^. 3P microscopy with 1700 nm excitation was first used in the intact mouse brain to image cortical and subcortical vasculature as well as RFP-labeled neurons throughout the visual cortex, cortical subplate, and hippocampus^12^. Since then, 1300 nm excitation of GCaMP6s has been used to record spontaneous neuronal activity throughout the cortical-subcortical volume^13^ and the sensitivity of these recordings and concomitant thermal and phototoxic changes have been characterized extensively^14^. Simultaneous dual color *in vivo* 3P microscopy remains challenging due to the large spectral separation in the optimal wavelength excitation windows for GFPs and RFPs, centered at 1300 and 1700 nm^3,12^, which currently necessitate independent dispersion compensation and custom optical elements with increased 1700 nm transmission.

Dual-color 3P microscopy of green and red fluorescent indicators was achieved in the fixed mouse brain using a novel two-stage optical parametric chirped-pulse amplifier seeded by a Ytterbium-doped fiber amplifier^15^ for 1300 nm and 1700 nm simultaneous output. However, issues associated with overlap of the two wavelengths at the focus and the requirement of higher powers (∼ 1 μJ) prohibit the application of this method to *in vivo* imaging in the mouse brain^14,16^. Alternatively, a recent report showed that single-wavelength 1300 nm excitation can be used with compatible combinations of fluorophores to achieve deep tissue simultaneous dual color imaging in the living mouse brain via higher-energy electronic excited states of the fluorophores^17^. However, a lack of information on the 3P excitation properties of commonly used fluorophores currently limits broad application of this approach. Here, we characterize the use of two RFPs with potentially enhanced excitation properties for *in vivo* single-source excitation 3P microscopy. These results will be important to. future studies utilizing this method to perform cell fate tracking, cell localization for holographic stimulation, and subcellular neuronal and glial imaging deep in the living mouse brain.

## 2. Methods

### 2.1. Animals

Animal experiments were conducted in accordance with protocols approved by the Animal Care and Use Committee at the University of Colorado Anschutz Medical Campus. Male and female mice used in these experiments were kept on a 14-h light–10-h dark schedule with ad libitum access to food and water and were housed with littermates. C57BL/6N *MOBP–EGFP* (MGI:4847238), *Olig2tm1(cre/Esr1*)Htak* (MGI:2183410), and *B6*.*Cg-Gt(ROSA)26Sortm9(CAG-tdTomato)Hze/J* (JAX #007909) were used for dual-color 3P imaging. Generation and genotyping of mice was performed as described previously^11^.

### 2.2. Custom three-photon microscope

A VIVO Multiphoton Open (3i) microscope, based on a Sutter Moveable Objective Microscope (MOM), was modified for three-photon imaging. The excitation source was a regenerative amplifier with 1030 nm center wavelength, 70 W average power, < 300 fs pulse duration, adjustable repetition rate up to 2 MHz (Spirit-1030-70, Spectra Physics), wavelength converted by a non-collinear optical parametric amplifier (NOPA-VIS-IR, Spectra Physics). The idler output of the NOPA is tunable in the 1200 - 2500 nm range. The laser was operated at a repetition rate of 1 MHz and the final output power from the idler was 0.8-1.1 W at 1300 nm. The power was modulated with a motorized half wave plate (KPRM1E/M - Ø1”, Thorlabs). Beam conditioning of the NOPA output consisted of a Glan-Thompson prism, and expansion and collimating lens relay (f1 = 75 mm, f2 = 100 mm, Newport), a 4x reflective beam expansion telescope (BE04R, Thorlabs), and a beam demagnifying telescope (f1 = 500 mm, f2 = 200 mm, Edmund Optics). The additional reflective beam expansion telescope and de-magnification were to condition the beam size for a deformable mirror that was held flat for this study. Group delay dispersion (GDD) compensation was achieved using a prism compressor system consisting of two 25mm SF10 prisms cut at Brewster’s angle (10NSF10, Newport), and a gold roof mirror (HRS1015-M01, Thorlabs). The beam was directed to the galvanometers (Cambridge Technologies) and through a scan lens (Thorlabs SL50-3P), tube lens and a 760 nm long-pass primary dichroic. The back aperture of a high-NA multiphoton objective (XLPLN25XWMP2, 25x/1.05 NA, Olympus) was ∼75% filled for 3P imaging. The fluorescent emission was separated from the excitation path by the long pass dichroic mirror and spectrally filtered (green channel = 525/50 nm, red channel = 620/60 nm), and detected by photomultiplier tubes (H10770PA-40, Hamamatsu). Electronic signal was amplified, low-pass filtered, and digitized. Data were acquired with SlideBook 2021 (Intelligent Imaging Innovations).

### 2.3. Two-photon microscopy

2P images in **Fig. S2** were generated using a wavelength tunable femtosecond oscillator (MaiTai-HP DeepSee, Spectra Physics) combined with the 3-photon excitation path with a 1030 nm long pass filter after the prism compressor and aligned to co-propagate with the 1300 nm light.

### 2.4. GDD Tuning and Pulse Measurements

To tune the prism compressor for optimizing 3P excitation, we performed time lapse imaging of a green fluorescent flat slide and adjusted the prism separation and insertion in real-time to maximize the fluorescent signal. We checked but did not need to make any changes in the compressor for maximum signal over multiple months of imaging. Pulse measurements were made using frequency-resolved optical gating (FROG) (FROGscan, Mesa Photonics). The FROG has an all-reflective optical path from its input to the BBO crystal used for second harmonic generation (SHG) to avoid adding any additional dispersion. For the same reason, a reflective objective 40x/.5 NA (LMM40X-UVV, ThorLabs) was used to collimate the beam after focusing through the objective, immersion water, and coverglass, followed by two silver mirrors to align into the FROG. This set up allowed us to characterize our pulse duration at the sample without additional dispersive elements that would change our pulse profile. We characterized the pulses for all wavelengths (**Fig. 1**, 1225 - 1360 nm). For the 2P excitation, we used the internal GDD tuning on the MaiTai-HP DeepSee to find the maximum signal intensity at each wavelength.

**Figure 1:**
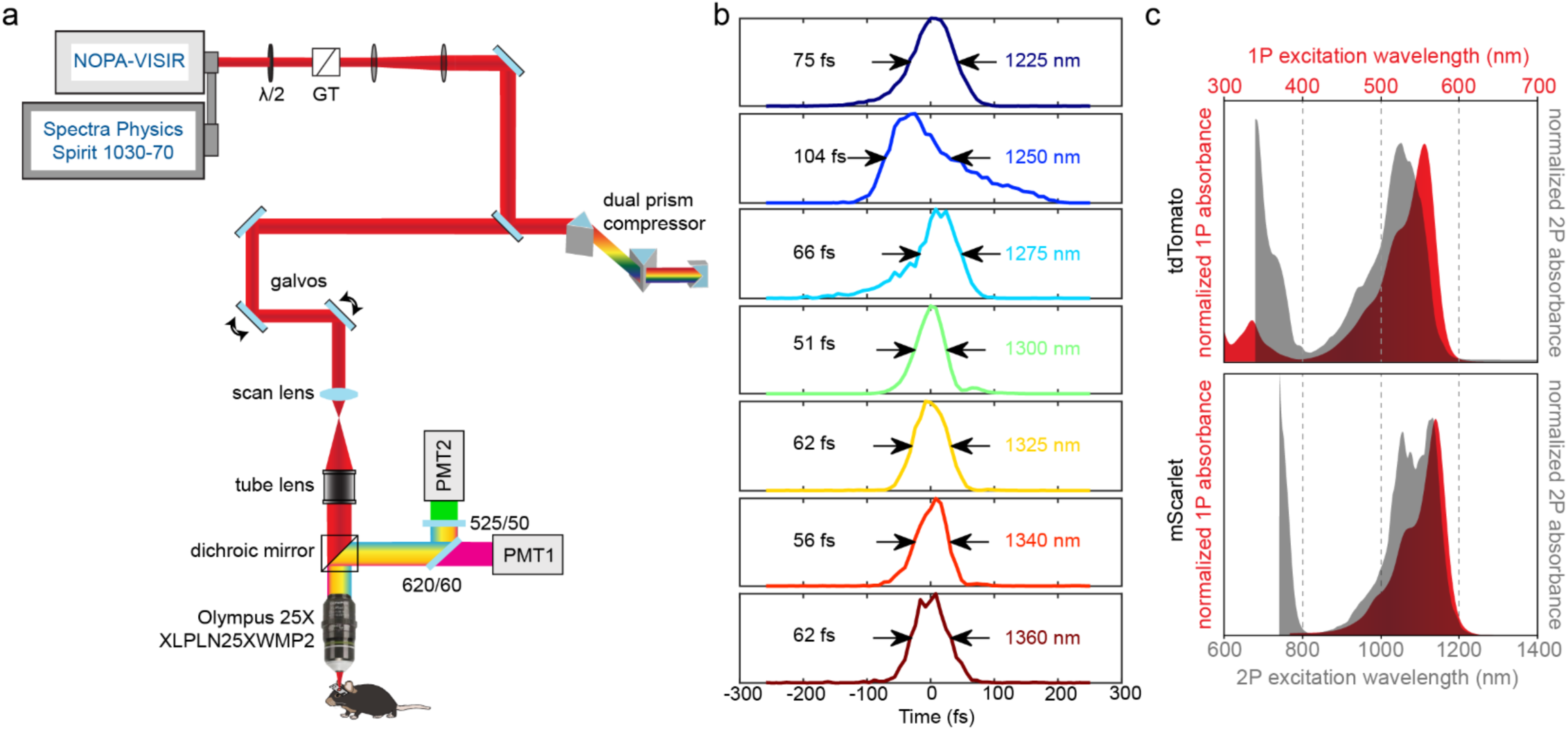
Custom three-photon microscope for dual color *in vivo* imaging. **a**) 3P light path and detection scheme for dual color imaging. **b**) Laser pulse temporal profile for wavelengths from 1200 – 1400 nm, using frequency-resolved optical gating. Pulses were characterized at the objective focus. Pulse widths indicated are the full width at half maximum (FWHM). **c**) 1P and 2P excitation curves for tdTomato and mScarlet, recreated from FPbase. Note the small tdTomato excitation peak in the UV and the short wavelength 2P excitation peaks at 750-800 nm.

### 2.5. Stereotaxic AAV injections

Six- to eight-week-old mice were anesthetized with Isoflurane inhalation (induction, 5%; maintenance, 1.5– 2.0%, mixed with 0.5 liter per min O_2_) and kept at 37 °C body temperature with a thermostat-controlled heating plate. A midline incision in the skin was made with fine surgical scissors to expose the skull. A small burr hole was made with a high-speed dental drill over the right forelimb primary motor cortex (1 mm anterior and 1.5 mm lateral to Bregma) and the deep layer of the thinned skull was removed with a curved 27-gauge needle. To label layer V/VI cortical neurons, we made two 500nL injections of AAV8-hSyn-mScarlet-WPRE^18^ (1.5 × 10^12^ vg/mL, Deisseroth lab, Stanford Viral Vector Core) at 1000 and 750 µm depths below the pial surface at a rate of 100 nL / minute. The Injection needle was left in place for 5 min. to prevent backflow and then removed slowly. The burr hole was filled with Vetbond (3M) and the skin incision was sutured using 5-0 Ethilon sutures.

### 2.6. *In vivo* multiphoton microscopy

Cranial windows were prepared as previously described^19^. Six- to ten-week-old mice were anesthetized and monitored as above. After removal of the skin over the right cerebral hemisphere, the skull was cleaned and a 2 × 2 mm region of skull centered over either a) the posterior parietal cortex (−1 to -3 mm posterior to bregma and 1 to 3 mm lateral), or b) the forelimb region of the primary motor cortex (0–2 mm anterior to bregma and 0.5–2.5 mm lateral) was removed using a high-speed dental drill. A piece of cover glass (VWR, No. 1) was placed in the craniotomy and sealed with Vetbond (3M) and then secured with dental cement (C&B Metabond). A 5 mg per kg dose of carprofen was subcutaneously administered before awakening and for three additional days for analgesia. For head stabilization, a custom metal plate with a 6 mm central hole was attached to the skull. The headbar was secured very close to the skull to enable deep 3P imaging without steric hinderance of the objective. Custom 3D-printed headbar holders (CU Anschutz Optogenetics and Neural Engineering Core) were attached to a 200 mm breadboard (Thorlabs) affixed to stacked dual-axis goniometers (Optosigma, +/- 15°, GOH-65B76R) such that the rotation center height was located at the headbar. Prior to imaging, the angle of the cranial window was adjusted to be perpendicular to the axis of the excitation beam using the goniometers. *In vivo* imaging measurements were taken either immediately following the surgery (acute preparation) or 2-3 weeks post-surgery (chronic preparation) as noted in the figures. During imaging sessions, mice were anesthetized with isoflurane and immobilized by attaching the head plate to the custom stage and temperature was monitored continuously as above. Images were acquired using Slidebook 2021 (3i) imaging software with 3 key modifications to standard laser point scanning systems for 3P imaging: 1) the laser was blanked during the galvanometer overscan to reduce the average power at the sample outside of the imaging FOV, 2) Slidebook 2021 was modified to allow for fine power control of the motorized half wave plate (+/- 0.1%), and 3) scanning was intermittently paused for 1 minute after every 3 minutes of continuous scanning to allow for heat dissipation^20^. Large 3D structural z stacks were acquired at 512×512 pixels, 2 µs dwell time, and frame averaging = 2, with a FOV of 385 × 385 µm with a 3 µm z-step. Z-stacks for optical measurements were acquired with the same settings without frame averaging and with a 0.5 µm z-step. The power after the objective was measured daily by centering the galvanometers and acquiring a point-scan power curve on a high-power microscope slide meter sensor head (S175C, Thorlabs) in immersion water. The maximum power at the sample was 1-55 mW (0-1100 mm depth).

### 2.7. Image Processing and Analysis

Image stacks were registered with StackReg (http://bigwww.epfl.ch/thevenaz/stackreg/) prior to analysis with ImageJ^21^. All images shown are either raw or filtered with a 0.5 µm median filter, as noted in the figure legends. Z-width of max projected images are also noted in the figures. Signal to Background Ratio (SBR) analysis was performed using ImageJ on 3-10 cells located ±5 microns from the indicated depth. For each wavelength and pulse energy measured, a z-stack was acquired, and an ROI was drawn over the cell body of interest, and the Plot Z-Axis Profile ImageJ command was used to find the z-plane of peak intensity. All measurements were made at this single plane image. For the peak signal, a line scan was made through the cell body and the five pixel values surrounding the maximum were averaged. For SBR measurements of mScarlet, the background fluorescence was measured within the area of an unlabeled neuron to avoid measuring virally labeled neurites as background fluorescence. Plotting and statistical analyses were performed in JMP 15 (SAS). SBR measurements were expressed as Mean (SD). Logarithmic pulse energy vs. signal plots (**Fig. 3c, Figs. S3-4**) were analyzed using linear regression (MSE, mean square error) and the slopes were expressed as Mean ± SEM. The pulse energy at the focal plane (z-depth) in **Figs. S2, 4** was calculated with the equation below, using a linear extrapolation of previously published experimental effective attenuation lengths (EAL) in the mouse neocortex^12,14,22^, where *P* = average power, *f* = laser repetition rate (1 MHz), and z = cortical depth:

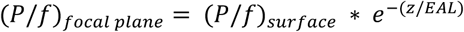

## 3. Results

### 3.1. Frequency-resolved optical gating pulse measurements

To overcome the fundamental depth limit of 2P imaging in the mouse brain^23^, we built a custom 3-photon microscope with a multichannel emission detection path (**Fig. 1a)**. With the ultimate goal of measuring the *in vivo* excitation properties of RFPs for simultaneous dual-color 3P microscopy, we first characterized our output pulses after the objective at a range of wavelengths using frequency resolved optical gating (FROG)^24^. Pulse characterization with FROG allows for the recovery of spectral intensity and phase using iterative spectrogram inversion algorithms and provides a ground-truth pulse width measurement^25,26^. We optimized the prism pulse compressor at 1300 nm to achieve 51 fs pulses and a pulse duration of 51-75 fs for 1225 to 1360 nm, except for 1250 nm, which was 104 fs (**Fig. 1b**). The raw spectrograms, retrieved spectral intensities, phases, and third-order fit GDD and TOD magnitudes are presented in **Figure S1**. The prism compressor was optimized at 1300 nm and maintained at the same settings for all measurements. The 1P and 2P excitation peaks for tdTomato are 554 and 1050 nm, while mScarlet is maximally excited at 569 nm and has a bimodal 2P excitation curve with peaks at 1065 and 1143 nm (**Fig. 1c**)^28, 29, 30, 31, 32,33^.

### 3.2. Nonlinear excitation properties of tdTomato in the living mouse brain at depth

We implanted chronic cranial windows over the posterior parietal cortex (PPC) of triple transgenic mice (*Olig2-CreER;RCL-tdTomato;MOBP-EGFP*,OTM) that express tdTomato in oligodendrocyte lineage cells and EGFP specifically in mature oligodendrocytes and myelin sheaths^11^. Single-excitation structural 3P microscopy of the brain volumes from OTM mice revealed single- and dual-labeled oligodendrocyte lineage cells throughout the cortical layers and subcortical white matter (**Fig. 2a-b**). To maintain constant average power at the brain surface, the laser power after the objective was measured at each wavelength and adjusted prior to imaging. We found that the pulse energies required for imaging were similar to those recently published for structural and functional neuronal imaging *in vivo*, and importantly, below the limits for heating- and nonlinear absorption-induced damage^14,16,34^. To determine the *in vivo* 3P excitation properties of tdTomato, we measured the fluorescent signal, background, and signal-to-background ratio (SBR) in layer 6b of the PPC (760 µm depth) of tdTomato at a range of wavelengths (1225-1360 nm) and constant average power at the surface (**Fig. 2c-g**). The peak signal increased at 1300 vs. 1225 and 1250 nm (mean (SD) = 1606.98 (499.02) vs. 879.66 (260.38), 386.79 (110.631), respectively), and plateaued through 1360 nm (**Fig. 2e**), consistent with a broad excitation range for tdTomato in the 1300 nm range. However, we found that the background decreased from 1300 to 1360 nm, resulting in an increased SBR in the 1325 - 1360 nm range when compared to 1300 nm (**Fig. 2g**, 21.91 (6.01), 22.39 (4.32), 30.74 (9.78) vs. 17.13 (5.32), respectively). To assess the contribution of 2P vs. 3P excitation processes to signal generation for each wavelength, we generated logarithmic signal vs. pulse energy plots at each wavelength for a range of pulse energies at the focal plane (2.3 – 3.8 nJ, **Fig. S2**). The slope of the logarithmic plot for tdTomato at 1225 nm was 2.171 ± 0.526, indicating a larger contribution of 2P than 3P excitation to the fluorescent emission at this wavelength and imaging depth. The log-plot slopes at 1300 and 1325 nm were 3.066 ± 0.390 and 2.874 ± 0.369, respectively, indicating that 3P excitation dominates in the 1300 nm range for tdTomato. These *in vivo* results are consistent with recently published 2P and 3P action cross sections of purified samples^17^. Our data indicate that the significant decrease in out-of-focus (2P) background with longer-wavelength 3P excitation of tdTomato drives the observed increase in the SBR of this fluorophore.

**Figure 2:**
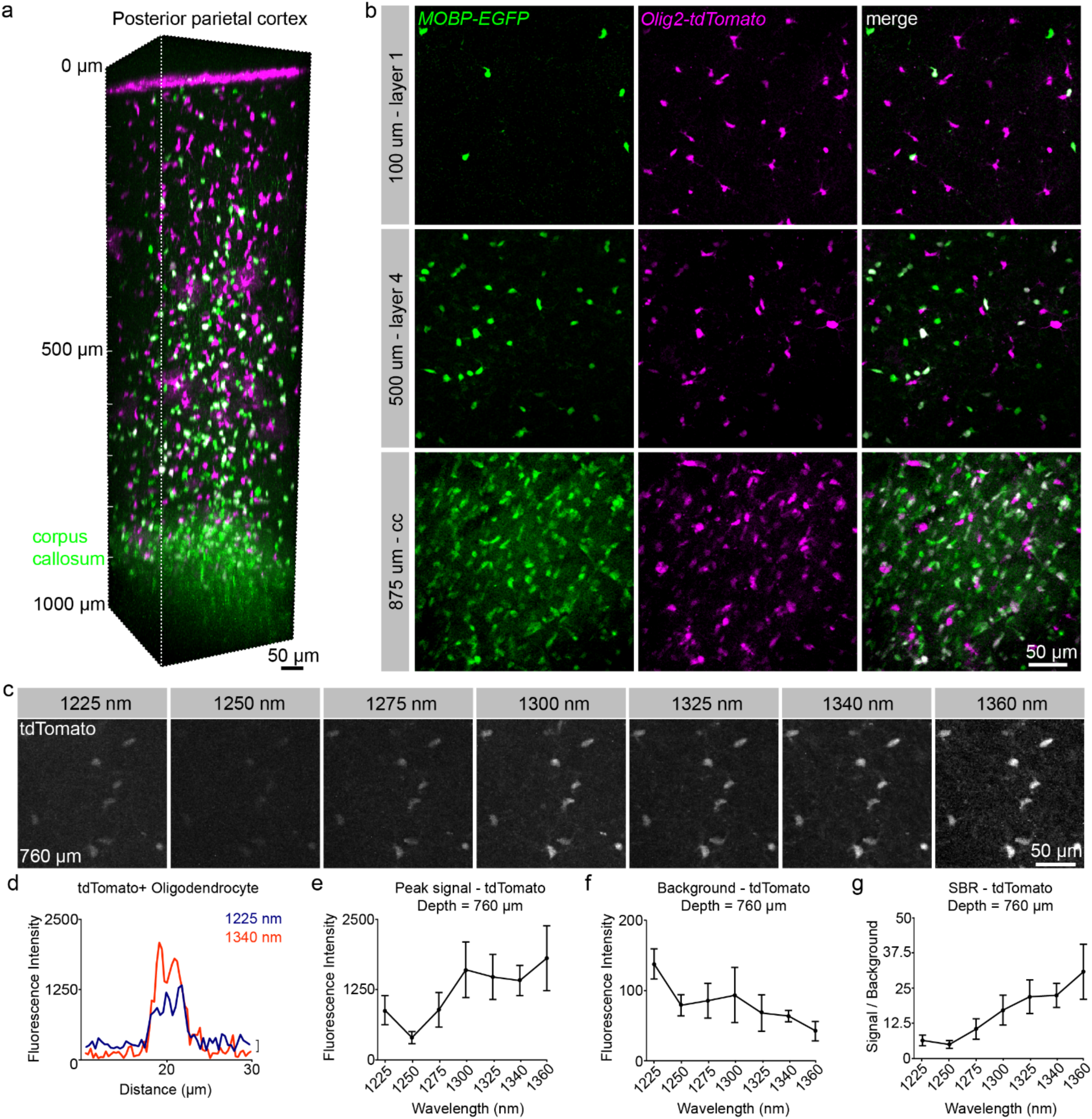
Simultaneous 3P excitation of EGFP and tdTomato in posterior parietal cortex (PPC). **a**) 3D image volume from a chronic implanted cranial window in an *Olig2-CreER; RCL-tdTomato; MOBP-EGFP* mouse at P60. **b**) Max projections of ∼50 µm volumes in cortical layers 1 (top), 4 (middle), and the corpus callosum (cc, bottom). Note the highly myelinated corpus callosum shows an increased density of dual-labeled mature oligodendrocyte cell bodies and strong *MOBP-EGFP* myelin signal compared to superficial cortex. **c**) Max projection images of 9 µm volumes at 760 µm depth and at a range of excitation wavelengths with average power = 45.2 mW. **d**) Example line scan plots through a single tdTomato-positive oligodendrocyte cell body at 1225 and 1340 nm show increased signal and decreased background with longer wavelength excitation. **e**) Max fluorescence signal of tdTomato-positive oligodendrocyte lineage cells. **f**) Background fluorescence in the tdTomato detection channel. **g**) Signal to background ratio vs. wavelength comparison of the tdTomato signal at 760 µm depth (**e-g**) Plots include measurements taken with average power = 45.2 mW after the objective and pulse energy = ∼2.5 - 3.6 nJ at the focus. Data represent n=10 cells from a triple transgenic mouse and are represented as the mean ± standard deviation. Due to low output power at 1360 nm, the data at this wavelength were acquired using 36 mW after the objective and then scaled by a value of 1.98 = (45.2 / 36.0)^3^.

### 3.3. Nonlinear excitation properties of mScarlet in the living mouse brain at depth

Historically, limitations in the maturation and quantum yield of monomeric RFPs have hampered their usage. Recently, a truly monomeric protein with high brightness and quantum yield, mScarlet, was developed^27^. This fluorophore has been included in a novel suite of viral tools used to express the highly sensitive, red-shifted opsin, ChRmine, to provide a red-shifted fluorescent reporter for structural imaging of targeted cells and subcellular structures^18^. To investigate the utility of mScarlet for dual color 3P microscopy, we injected AAV8-hSyn-mScarlet-WPRE into the deep layers of the primary motor cortex to label motor output neurons. These injections were performed in *MOBP-EGFP* mice, which express EGFP in myelinating oligodendrocytes, to allow visualization of neuron-glia interactions. Single-excitation structural 3P microscopy of mScarlet-injected *MOBP-EGFP* mice permitted the simultaneous visualization of excitatory neurons and mature oligodendrocytes at depths up to 1100 µm in forelimb motor cortex (**Fig. 3a**). Large diameter layer 5 / 6 neuronal cell bodies were labeled at depths greater than 800 µm and mScarlet-positive processes were labeled throughout the cortical volume (**Fig. 3a-b**). To characterize the 2P excitation profile of mScarlet, we measured the *in vivo* 2P SBR of mScarlet at a range of wavelengths centered at 950 nm at a depth of 400 µm from the brain surface. We found that the *in vivo* SBR of the two fluorophores intersected at ∼960 nm suggesting that this may be an ideal wavelength for 2P dual color imaging of EGFP and mScarlet (**Fig. S3**). However, we found acceptable dual-color cellular signal was achieved from 920 - 1000 nm, highlighting the broad applicability of this fluorophore combination at wavelengths with sufficient output power from standard multiphoton lasers. These *in vivo* 2P excitation data build upon recently published mScarlet cross section results in solution showing a bimodal excitation curve with an initial peak near 1060 nm (**Fig. 1c**)^29^. Next, we measured the 3P SBR of mScarlet in the deep primary motor cortex (920 µm below the brain surface) at a range of excitation wavelengths and constant average power at the surface. We found that mScarlet showed a large signal increase at 1225 nm yet had a high out-of-focus background that substantially reduced the SBR at this wavelength (**Fig. 3c-f**), likely due to the predominance of 2P over 3P excitation at shorter wavelengths. While the mScarlet signal initially decreased from 1250 – 1300, it subsequently increased from 1325 to 1360 nm (**Fig. 3d**). Unexpectedly, we found that the mScarlet background plateaued from 1300 - 1360 nm, despite choosing ROIs within unlabeled neuronal shadows. However, these *in vivo* measurements may still overestimate the background at longer wavelengths due to strong diffuse labeling in the AAV-injected mice (**Fig. 3c, e**). The mean mScarlet SBR plateaued from 1340 - 1360 nm, which may represent an excitation peak for higher order excitation in the 1300 nm range (**Fig. 3f**). To determine the contributions of 2P versus 3P excitation we generated logarithmic signal vs. pulse energy plots at each wavelength for mScarlet as above (**Fig. S4**). We found that the slope of the logarithmic mScarlet plot at 1225 nm was 1.885 ± 0.896, reflecting the strong contribution of 2P excitation at this wavelength (**Fig. S4, Fig. 3c-d**). In contrast, we found that the slope at 1340 nm was 2.915±2.05, confirming the predominant contribution of 3P excitation at wavelengths greater than 1325 nm for mScarlet (**Fig. S4**). Together, our data suggest that mScarlet is an attractive RFP with enhanced 3P cross section in the 1300 nm range for use in single excitation source dual color *in vivo* 3P imaging experiments.

**Figure 3:**
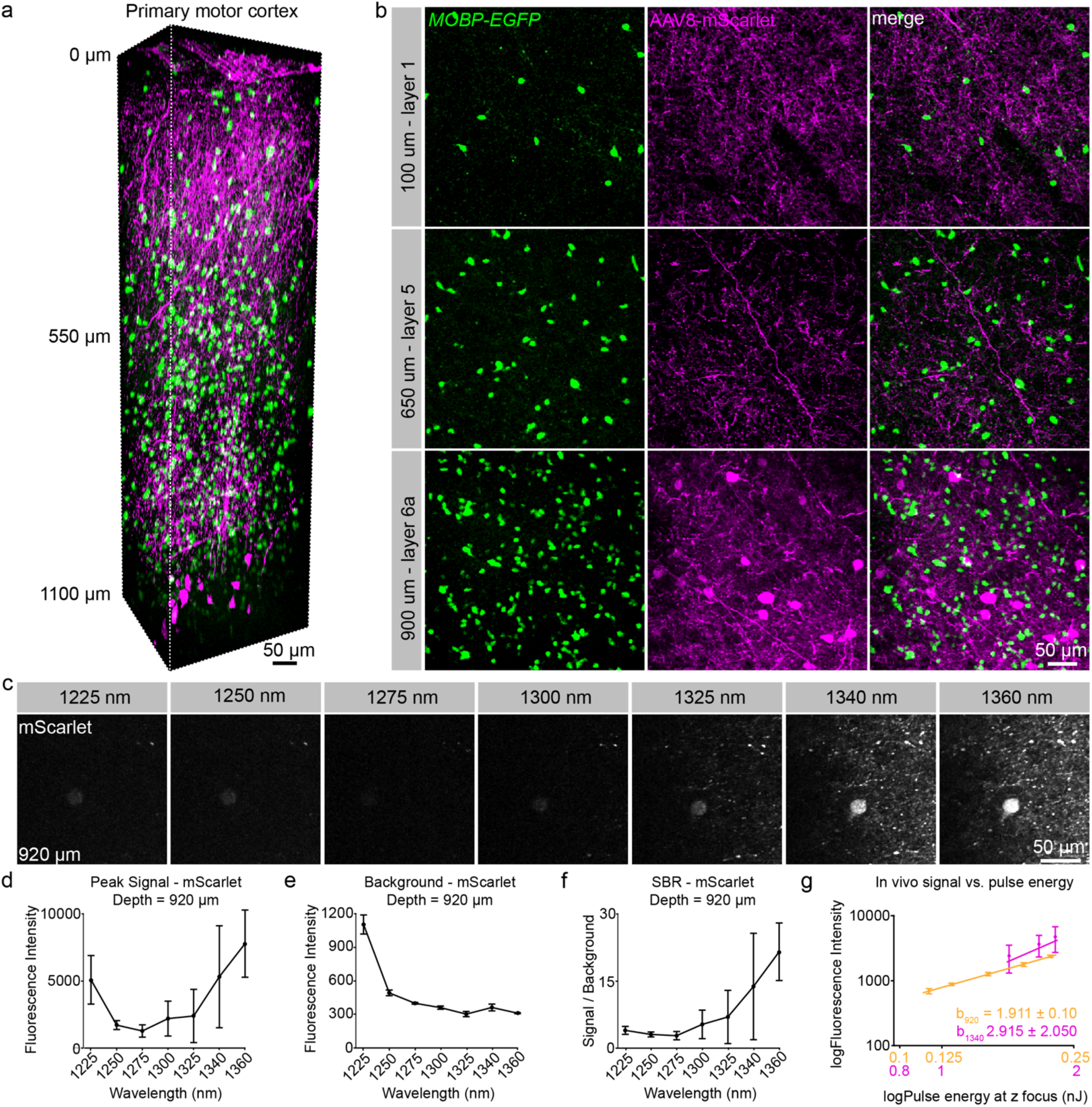
Simultaneous 3P excitation of EGFP and mScarlet in primary motor cortex. **a**) 3D image volume from an acutely implanted cranial window in an *MOBP-EGFP* mouse at P65 that was injected with AAV8-hsyn-mScarlet virus at 1000 and 750 µm depths in the primary motor cortex. Note large mScarlet-positive layer 5/6 motor output neurons are labeled at the bottom of the image volume and neuronal processes of these cells are labeled throughout the volume. **b**) Max projection images of ∼30 µm volumes in cortical layers 1 (top), 5 (middle), and 6a (bottom). **c**) Max projection images of 9 µm z-width at 920 µm depth show differences in signal and background fluorescence across a range of excitation wavelengths. Note the increased signal and background at 1225 nm. **d**) Max fluorescence signal of mScarlet-positive neurons for the range of wavelengths. **e**) Background fluorescence measurements in the mScarlet channel. **f**) Signal to background ratio vs. wavelength comparison of mScarlet signal at a depth of 920 µm from the brain surface. **g**) log-log signal vs. pulse energy plots illustrate the contribution of 2- and 3P excitation to the fluorescence intensity for mScarlet at 920 (orange) and 1340 nm (magenta). The steeper slope of the 1340 nm line plots indicates a greater contribution of 3P excitation to the signal at the longer wavelength (theoretical 3P/2P ratio = 1.5). **d-g**) plots include measurements taken with average power = 47.7 mW after the objective and pulse energy = ∼1.3 - 2 nJ at the focus. Data represent n= 3 labeled neurons from a virally injected transgenic mouse and are represented as the mean ± standard deviation. Due to low output power at 1360 nm, the data at this wavelength were acquired using 38.5 mW after the objective and then scaled by a value of 1.90 = (47.7 / 38.5)^3^.

### 3.4. Modeling of the excitation probability and brain heating for tdTomato in the 1300-1650 nm range

Simultaneous dual-color 3P excitation in the 1300 nm range may require excitation of fluorophores at sub-optimal excitation wavelengths, resulting in increased excitation power and heating. To model the probability of excitation for tdTomato we used 3P action cross section data^17^ and infrared water absorption values^35^ in the equation below^14^:

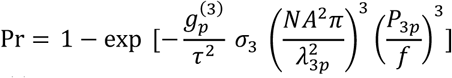

Where 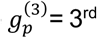 order temporal coherence factor, t = pulse duration (s), *σ*_3_ = 3P cross section, NA = objective numerical aperture, λ = excitation wavelength, P_3p_ = photons / s at the focal plane, and *f* = laser repetition rate. The modeled probabilities of excitation for 1300, 1340, and 1650 nm are presented in **Fig. S5a** and the power at the surface values required for 10% excitation probability per pulse are presented in **Fig. S5b**. With 30mW at the surface and z = 800 µm, the excitation probability per pulse is .07, .11, .26 for 1300, 1340, and 1650 nm. Therefore, 10% excitation probability is achieved with 34.5, 29, and 21 mW average power at the surface for 1300, 1340, and 1650 nm, respectively, for the specific parameters of our microscope. The water absorption coefficient at 1300 nm is ∼22% of that at 1650 nm (.136 vs .616 mm^-1^)^35^. Because heating scales nearly linearly with input power^14^ optimal excitation vs. heating is achieved in the 1300 nm range. Additionally, the 3P action cross section for tdTomato is slightly greater at 1320 nm than 1340 nm (3.3 vs. 3.13 *10^−82^ cm^6^(s/photon)^2^)^17^, which is in agreement with our *in vivo* SBR values (**Fig. 2g**). Therefore, ∼1320 nm is the optimal excitation wavelength for single-wavelength 3P excitation of EGFP and tdTomato. Our *in vivo* SBR measurements for mScarlet (**Fig. 3f**) suggest that the optimal excitation wavelength will be slightly red-shifted from tdTomato, however future 3P action cross section experiments for mScarlet, combined with empirical fine-tuning of excitation parameters will be essential in determining these values for *in vivo* imaging.

## 4. Discussion

### 4.1. tdTomato and mScarlet have enhanced three-photon cross sections in the 1250 – 1350 nm range

A recent study showed that while excitation in the 1600 – 1700 nm range resulted in appreciable 3P excitation to the lowest energy state, certain RFPs have an order of magnitude larger 3P action cross section in the 1260 – 1360 nm range^17^. Here, we took advantage of these 3P excitation properties to characterize the *in vivo* behavior of a classical tandem dimer RFP (tdTomato), as well as a recently developed monomeric RFP (mScarlet), for use in simultaneous dual-color 3-photon imaging studies. We found that both tdTomato and mScarlet have favorable 3-photon excitation properties in the 1300 – 1340 nm range and we empirically defined the optimal excitation wavelengths for the best SBR at depth, which will be applicable to future dual-color *in vivo* imaging experiments. Similar to Hontani and colleagues^17^, we found that tdTomato exhibited a broad 3P excitation curve that increases significantly from 1250 – 1300 nm and plateaus through 1360 nm. We found that the improvement in the signal to background ratio at the longer wavelengths (1300 – 1360 nm) was largely due to decreased out-of-focus background generation at these wavelengths. In contrast, the signal vs. wavelength curve for mScarlet was slightly red-shifted, as the peak signal increased significantly from 1325 – 1360 nm wavelengths, however, mScarlet also showed exceptional signal to background in this range. These results suggest that both of these red fluorophores have advantageous excitation properties in the short 3P wavelength range (1300 – 1340 nm). Future characterization of additional red fluorophores in the 1300 – 1700 nm range will be useful to direct the application of both structural and functional *in vivo* dual-color 3P imaging.

### 4.2. 1300 – 1360 nm excitation of RFPs likely represents excitation to a higher energy molecular state

Certain red fluorescent proteins like Texas Red, tdTomato, and mScarlet exhibit short-wavelength one-photon absorption bands (∼400 – 450 nm) that represent excitation to higher-energy electronic states. Multiphoton excitation wavelengths used with classically employed fluorescent indicators likely induce excitation to the lowest-energy excited state (absorption peak), which has empirically generated optimal signal to noise in 2P experiments (e.g., 1040 nm excitation of tdTomato). However, our current findings suggest that 3P excitation is enhanced in these lower-wavelength absorption bands and has considerable implications for the design and implementation of future dual-color 3P studies. In addition, the ∼1260 – 1340 nm range of 3P excitation wavelengths overlaps with the long-wavelength tail of the 2P absorption cross section for most RFPs, therefore, signal detected at wavelengths less than ∼1300 nm represents a mix of 2P and 3P excitation. We found that the mScarlet signal detected at 1225 nm was greater than in the 1250 – 1325 range, likely due to the low energy tail in the mScarlet 2P absorption cross section curve. However, consistent with 2P excitation dominating at this wavelength, we measured high out-of-focus background generation at 1225 nm that severely decreases contrast and SBR at increased depths.

### 4.3. Complete characterization of laser pulses for three-photon excitation in the 1200-1360 nm wavelength range

We used frequency resolved optical gating (FROG) to measure the electric field of our laser pulses at the different wavelengths used for 3P microscopy. Previously, autocorrelation measurements in the 1200 – 1400 nm range have been presented^12,17^. FROG measurements have an advantage over autocorrelation in that both the intensity and phase of the pulse can be retrieved from the spectrograms and is useful for pulses with complex temporal profiles^25^. The retrieval of the electric field is done without prior assumptions of pulse shape, which are required for estimating pulse widths from autocorrelation measurements. Calculation of time-bandwidth product and the spectral phase can determine whether the pulse is optimally temporally compressed at the sample. While there are a variety of FROG instrument geometeries^25^, we used second harmonic generation (SHG)-FROG. One limitation of SHG-FROG is that the direction of time of the pulse is not captured in the measurement^37^. This makes time-reversed traces in **Fig. 1** equally likely and leads to ambiguity in the sign of the spectral phase. We measured an asymmetric pulse profile and longer pulse duration for 1250 and 1275 nm center wavelengths when tuning the output from the NOPA laser. Full characterization and correction for dispersion to allow the shortest pulse at the sample is especially important for 3P microscopy where the fluorescent signal, I_S_, is highly dependent on the pulse duration, t, by the relation, 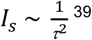.

### 4.4. Signal to background ratio dependence on pulse duration

When tuning the wavelength from 1225 to 1360 nm, the excitation of tdTomato changes from a predominately 2P process, to a mixture of 2P/3P, to a 3P process. Interestingly, when there is a mixture of 2P/3P, the SBR shows strong dependence on the pulse duration (∼1/t). However, for a fully 2P process or fully 3P process where the background is above the detector noise level, there is a weaker dependence of SBR on pulse duration (see Ref. 38, Fig. S4 and Ref. 39, Fig. S5).

For mixed 2P/3P excitation, we can approximate the SBR as^17^:

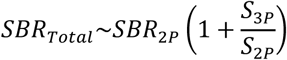

The ratio of 3P signal versus 2P is affected by both the pulse duration (FWHM), t, and the 2^nd^ and 3^rd^ order temporal coherence of the pulse (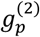 and 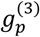)^17^ in the form:

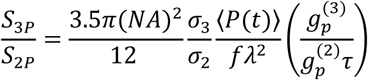

where *σ*_2_ and *σ*_3_ are the two- and three-photon cross sections, ⟨*P*(*t*)⟩ is the time-averaged excitation photon flux (photons/s), NA is the numerical aperture, f is the laser repetition rate, and *λ* is the excitation wavelength. Therefore we expect a lower SBR at 1250 and 1275 nm due to the longer pulse durations and asymmetric pulse shapes at those wavelengths.

As a further illustration, we theoretically calculated the SBR_Total_/SBR_2P_ for tdTomato as a function of pulse duration using the 2P and 3P cross sections measured at 1260 nm excitation and the following parameters: 2 nJ pulse energy, 1 MHz repetition rate, 0.8 NA, 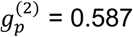 and 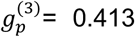 for hyperbolic-secant-squared pulse. Results are shown in **Figure S6**. As can be seen, the value for a 100 fs pulse is 53.7% lower than for a 50 fs pulse. The 3p excitation cross-section versus wavelength for mScarlet has not been measured so it is uncertain over which wavelength range the 2P/3P transition occurs.

### 4.5. Advantages of single wavelength dual-color 3P imaging

Optical engineering of microscopes capable of simultaneous 1300 and 1700 nm 3P excitation presents multiple difficulties. Current commercially available optical elements (e.g., Pockels cells) are not commonly tested in the 1700 nm range and achieving adequate transmission of the excitation light out the objective may require additional purchases and/or custom modifications. Additionally, 1700 nm light requires independent pulse dispersion optimization, and for simultaneous imaging, multiple excitation sources must be mixed and aligned for each imaging session. The results presented here suggest that for certain applications of structural dual color 3P excitation, specific fluorophore combinations can be used to exploit the enhanced 3P cross sections of RFPs in the 1300 nm range without the need for a second 1700 nm excitation path. Still, the development of simultaneous 1300/1700 nm systems will be important for future studies using dynamic optochemical indicators and/or optical stimulation deep in the brain. Furthermore, it was reported that unlike tdTomato and mScarlet, mCherry exhibits a 7-fold decreased 3P cross section in the 1300 nm range compared to 1650 nm^17^ and thus may require a 1300/1700 nm system to be optimally employed as a fluorescent reporter for dual-color 3P imaging.

### 4.6. Scattering and absorption in the 1300 – 1400 nm range

The effective attenuation length (EAL) for 3P microscopy is defined as the depth at which the unscattered excitation light is reduced by 1/e^3^. In the mouse brain, both 2P and 3P EAL are determined by a combination of scattering and water absorption. 2P and 3P EALs in different regions of the mouse brain have been characterized extensively, both *in vivo* and *in silico*^12,14,20,22,23,27,40,41^. Therefore, considerations such as the scattering and water absorption, as well as the output power from current NOPA configurations should be taken into account for dual-color 3P imaging with a single excitation source in the 1300 nm range. In this study, we did not achieve sufficient output power at 1 MHz and wavelengths greater than 1340 nm to allow for practical deep *in vivo* imaging, however, excitation in the 1325 – 1340 nm range provided exceptional SBR for both tdTomato and mScarlet. Because water absorption increases exponentially between 1340 and 1400 nm, our results are encouraging in that a narrow excitation window (1320 – 1340 nm) may be used to excite a broad range of green and red fluorescent reporter combinations with a single wavelength. Future work includes empirically testing a suite of fluorescent reporters for use in multicolor 3P experiments, using emission path engineering to increase the number of detection channels, and defining 3P activation properties of opsins and other dynamic indicators.

## Acknowledgments

We thank Dr. Chris Xu for his invaluable advice in setting up the 3P microscope at the University of Colorado Anschutz Medical Campus; Dr. Karl Deisseroth for providing us with the AAV8-hSyn-mScarlet-WPRE virus; Anthony Chavez for technical and veterinary assistance; Andrew Scallon and the CU Anschutz Optogenetics and Neural Engineering Core (P30NS048154) for 3D printing and stage design; members of the Hughes and Welle labs for discussions.

## Contributions

E.G.H, E.A.G., and M.A.T. conceived the project. G.L.F and E.A.G. built the microscope. M.A.T. conducted experiments and generated Figs. 1a, 2, 3, S1a, S3, S4, and S5. G.L.F. designed and contributed Figs. 1b, S1b, and S2. M.E.S. analyzed data in Figs. 2, 3, S3, and S5. B.N.O. and K.K. developed the software and helped with microscope setup. E.G.H., E.A.G., and D.R. supervised all experiments. M.A.T. and E.G.H. wrote the manuscript with input and editing from all other authors.

## Data and materials availability

All data that support the findings, tools, and reagents will be shared on an unrestricted basis; requests should be directed to the corresponding authors.

## List of Supplementary Materials

Supplementary Figures 1 to 6

