## Supplemental Materials for "Characterization of red fluorescent reporters for dual-color *in vivo* three-photon microscopy"

a

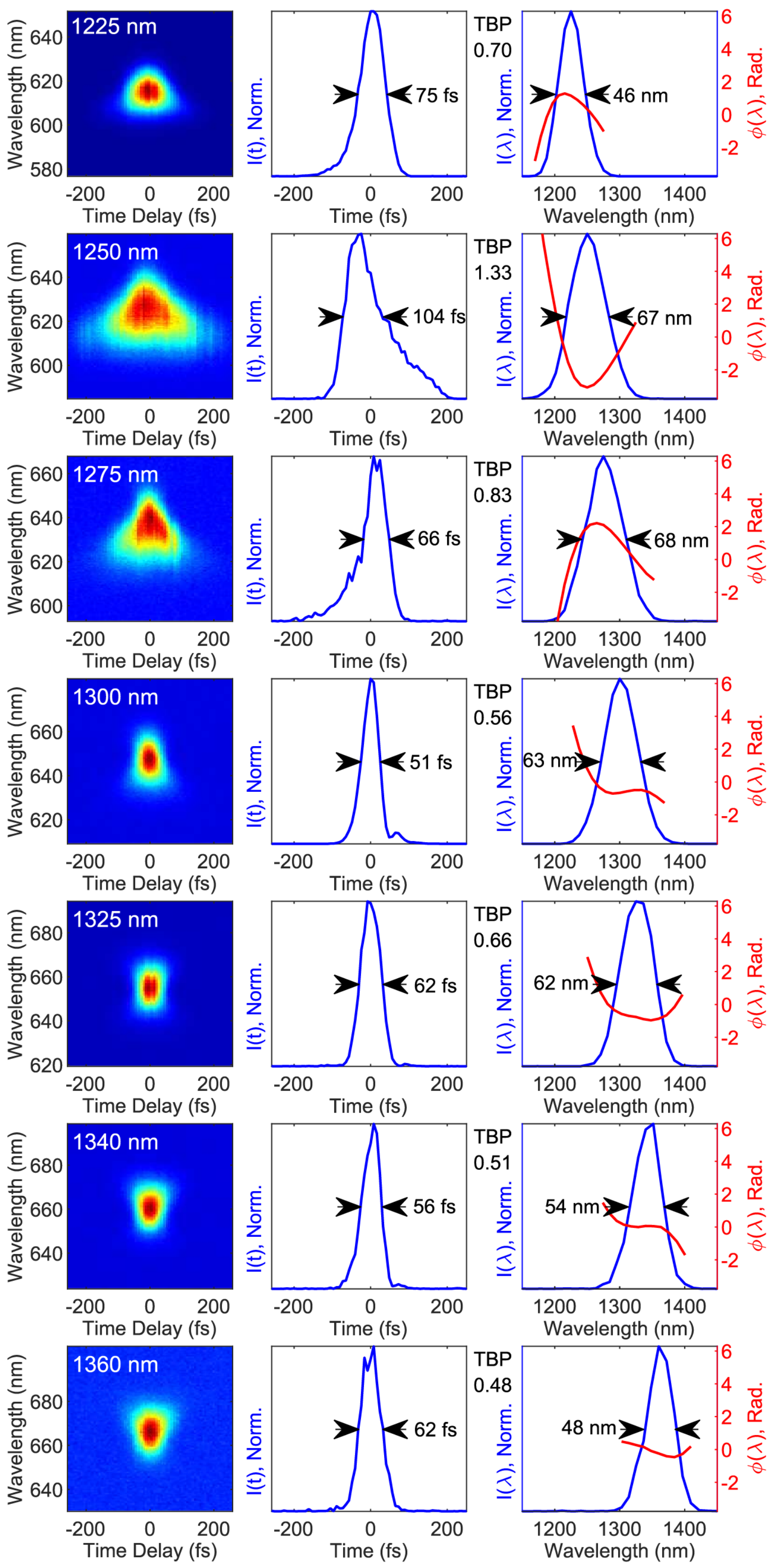


b

| λ (nm) | GDD (fs$^2$) | TOD (fs$^3$) |
|---|---|---|
| 1225 | 634 | 4302 |
| 1250 | 859 | 2817 |
| 1275 | 609 | 4196 |
| 1300 | 191 | 4940 |
| 1325 | 386 | 5895 |
| 1340 | 249 | 4532 |

**Supplementary Figure 1: Measured FROGscan spectrograms and retrieved temporal and spectral profiles. a, left**) Measured FROGscan spectrograms at each investigated wavelength. a, **middle**) Retrieved temporal intensity I(t), as in Fig. 1. (a, **right**) Retrieved spectral intensity, I(λ), with overlaid spectral phase, φ(λ), in radians (red) with time-bandwidth product (TBP) annotated between the traces. The larger phases on the pulses at 1250 & 1275 nm led to broader temporal pulses at these wavelengths and larger TBP. **b**, table showing the magnitude of group delay dispersion (GDD) and third-order dispersion (TOD) required to correct for the chirp of the pulse.

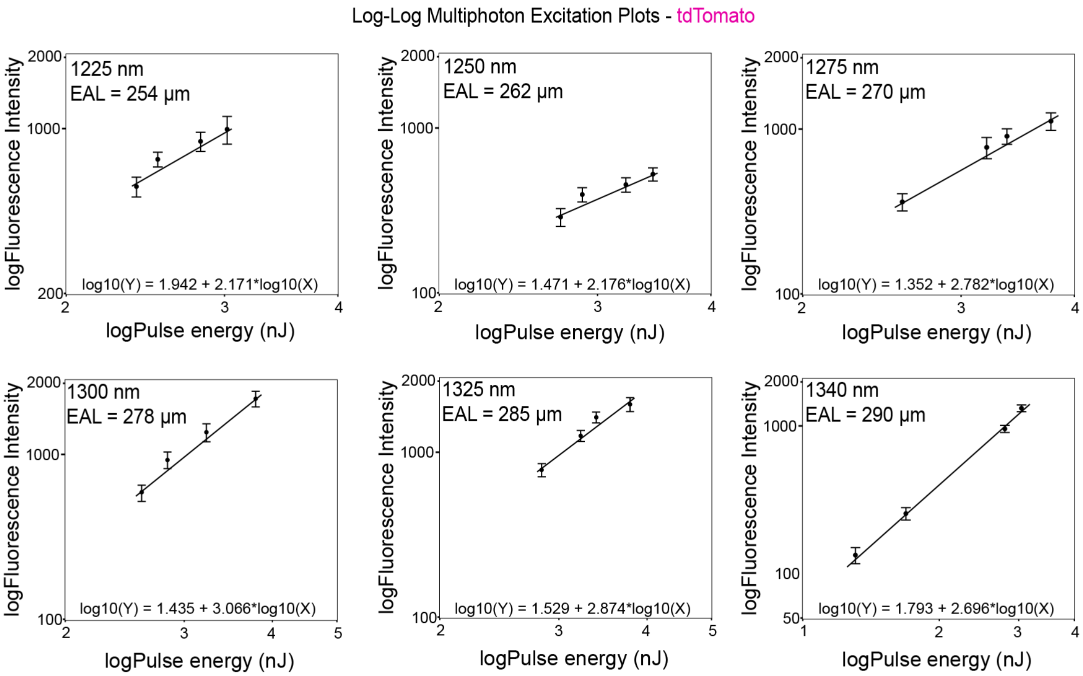


| Wavelength (nm) | Slope |
|---|---|
| 1225 | 2.171 ± 0.526 |
| 1250 | 2.176 ± 0.524 |
| 1275 | 2.782 ± 0.382 |
| 1300 | 3.066 ± 0.390 |
| 1325 | 2.874 ± 0.369 |
| 1340 | 2.696 ± 0.132 |

**Supplementary Figure 2: Log-Log pulse energy vs. signal plots for tdTomato at 760 µm below the brain surface. a)** individual logarithmic plots of pulse energy vs. *in vivo* fluorescent signal at a range of 3P wavelengths. **b)** Table of slope values for the excitation power ramp lines shown in (a). The steeper slope at higher wavelengths indicates a greater contribution of 3P vs. 2P excitation.

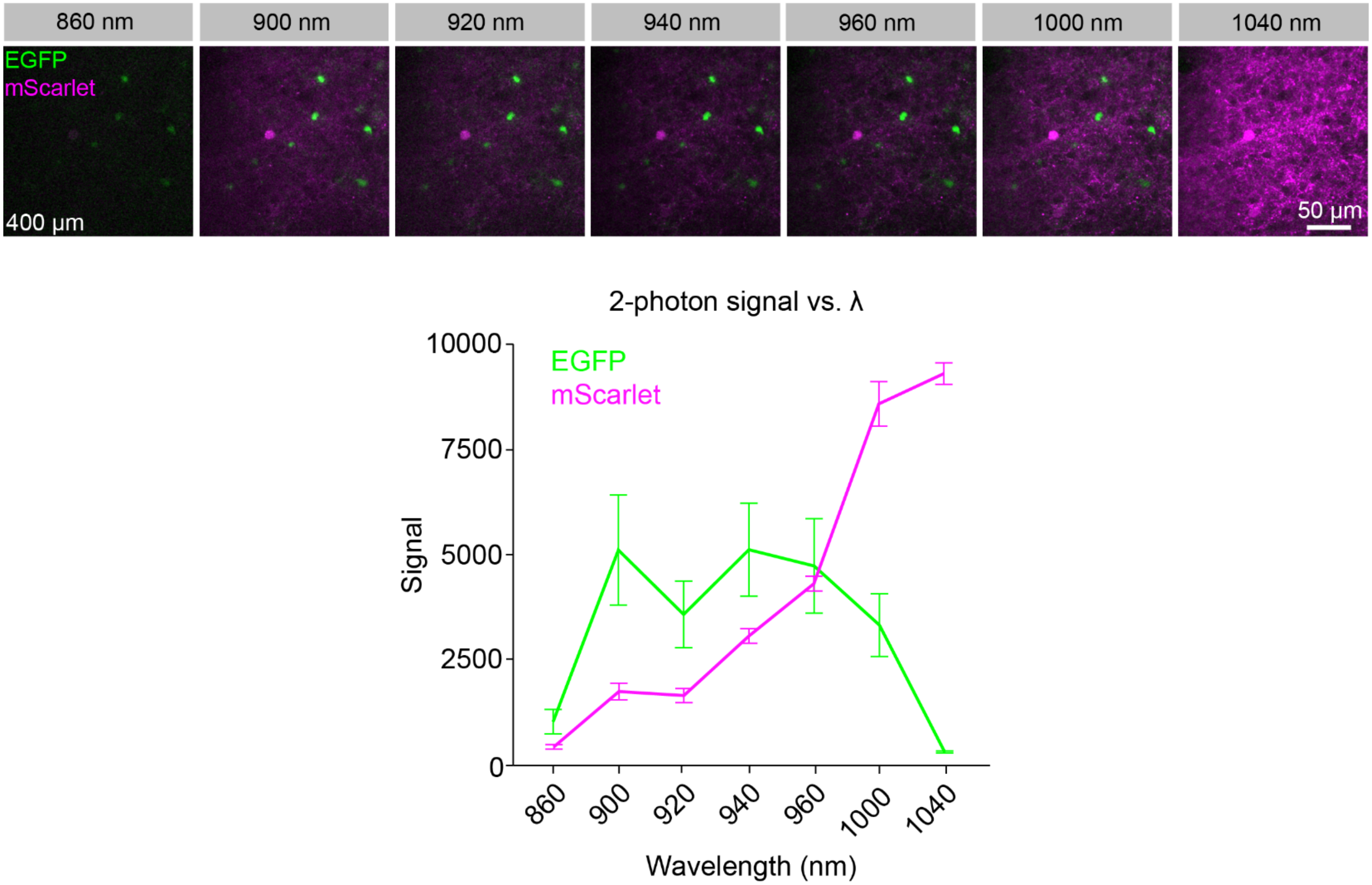


**Supplementary Figure 3: Two-photon *in vivo* imaging of EGFP and mScarlet at a depth of 400 µm from the pial surface.** Single plane images from a chronic implanted cranial window in an AAV8-hSyn-mScarlet injected *MOBP-EGFP* mouse with varying excitation wavelength (860 – 1040 nm) and quantification of *in vivo* signal generation compared to excitation wavelength. The pulse energy at the focus was .25 nJ at 920 nm, the average power at the surface was kept constant, and the pulse width was adjusted to maximum signal at each wavelength.

a

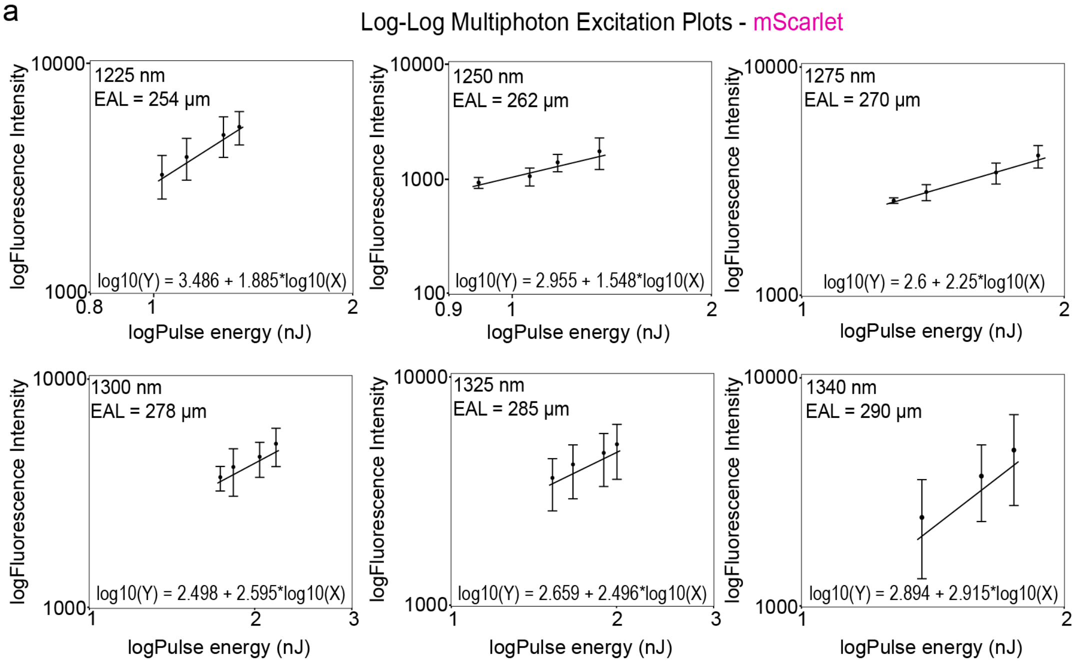


b

| Wavelength (nm) | Slope |
|---|---|
| 1225 | 1.885 ± 0.896 |
| 1250 | 1.548 ± 0.714 |
| 1275 | 2.250 ± 0.552 |
| 1300 | 2.595 ± 1.688 |
| 1325 | 2.496 ± 2.209 |
| 1340 | 2.915 ± 2.050 |

**Supplementary Figure 4: Log-Log pulse energy vs. signal plots for mScarlet at 920 μm below the brain surface. a**) Individual logarithmic plots of pulse energy vs. *in vivo* fluorescent signal at a range of 3P wavelengths. **b**) Table of slope values for the excitation power ramp lines shown in (**a**).

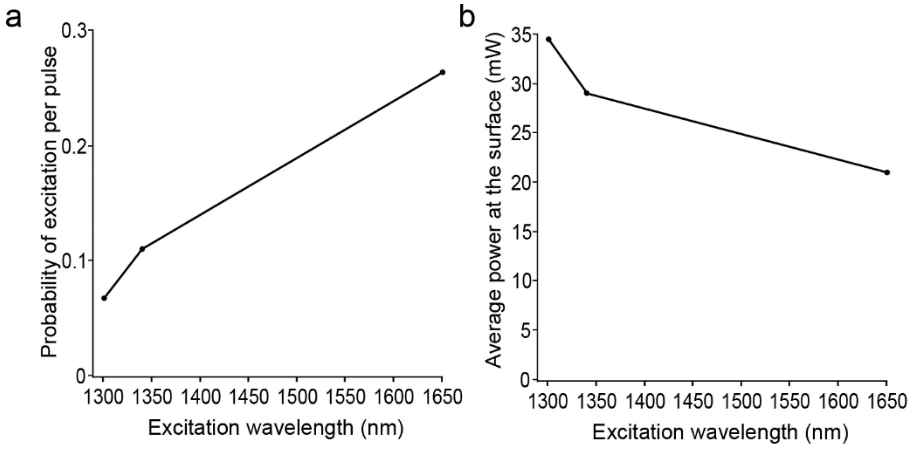


**Supplementary Figure 5: Computational modeling of 3P excitation probability and average power for tdTomato at 1300-1360 nm. a**) Probability of excitation per pulse at z depth = 800 µm with 30 mW average power at the surface. **b**) Average power at the surface required for 10% probability of excitation per pulse for 1300, 1340, and 1650 nm. Repetition rate = 1 MHz, objective NA = 0.8 (75% filled 1.05 NA Olympus objective), $EAL_{1300}$ = 278, $EAL_{1340}$ = 293, $EAL_{1650}$ = 350 µm, pulse duration = 60 fs.

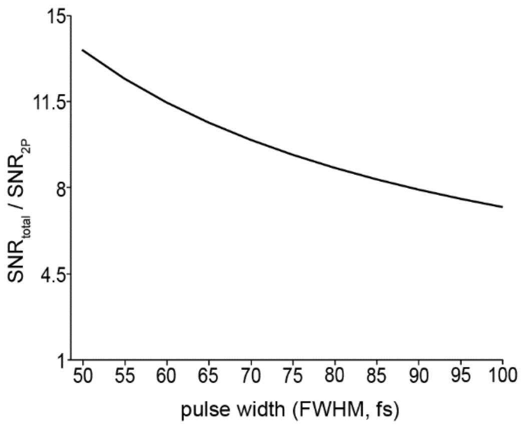


**Supplementary Figure 6: Theoretical calculation of the signal to noise enhancement ($SBR_{total}/SBR_{2p}$) for different pulse durations for tdTomato.** Parameters = 2 nJ pulse energy, 1 MHz repetition rate, 0.8 NA, and excitation wavelength of 1260 nm. Pulses were assumed to be hyperbolic-secant-squared. Exact values for the two- and three-photon cross sections were obtained from Yusaku Hontani and Chris Xu, Ref. 17.
